# Structural characterization of the essential cell division protein FtsE and its interaction with FtsX in *Streptococcus pneumoniae*

**DOI:** 10.1101/2020.06.08.141481

**Authors:** Martin Alcorlo, Daniel Straume, Joe Lutkenhaus, Leiv Sigve Håvarstein, Juan A. Hermoso

## Abstract

FtsEX is a membrane complex widely conserved across diverse bacterial genera and involved in critical processes such as recruitment of division proteins and in spatial and temporal regulation of muralytic activity during cell division or sporulation. FtsEX is a member of the ABC transporter superfamily, where FtsX is an integral membrane protein and FtsE is an ATPase, required for mechanotransmission of the signal from the cytosol through the membrane, to regulate the activity of cell-wall hydrolases in the periplasm. Both proteins are essential in the major human respiratory pathogenic bacterium, *Streptococcus pneumoniae* and interact with the modular peptidoglycan hydrolase PcsB at the septum. Here, we report the high-resolution structures of pneumococcal FtsE in complex with different nucleotides. Structural analysis reveals that FtsE contains all the conserved structural motifs associated with ATPase activity, and allowed interpretation of the *in vivo* dimeric arrangement in both ADP and ATP states. Interestingly, three specific FtsE regions were identified with high structural plasticity that shape the cavity in which the cytosolic region of FtsX would be inserted. The residues corresponding to the FtsX coupling helix, responsible for FtsE contact, were identified and validated by *in vivo* mutagenesis studies showing that this interaction is essential for cell growth and proper morphology.

**IMPORTANCE:** Bacterial cell division is a central process that requires exquisite orchestration of both the cell wall biosynthetic and lytic machineries. The essential membrane complex FtsEX, widely conserved across bacteria, play a central role by recruiting proteins to the divisome apparatus and by regulating periplasmic muralytic activity from the cytosol. FtsEX is a member of the Type VII family of the ABC-superfamily but instead transporter, couple ATP hydrolysis by FtsE to mechanically transduce a conformational signal to activate PG hydrolases. So far, no structural information is available for FtsE. Here we provide the structural characterization of FtsE confirming its ATPase nature and revealing regions with high structural plasticity key for FtsX binding. The complementary region in FtsX has been also identified and validated *in vivo*. Our results provide evidences on how difference between ATP and ADP states in FtsE would dramatically alter FtsEX interaction with PG hydrolase PcsB in pneumococcal division.

## INTRODUCTION

In order to divide, bacteria must synthesize a new septal cell wall in which the main component is peptidoglycan (PG). This newly synthetized PG must be split down the middle by one or more murein hydrolases to separate the resulting daughter cells. These biosynthetic and hydrolytic processes must be tightly coordinated and regulated to warrant bacterial survival. Although the regulation of these processes for PG hydrolases are not well understood, a principle that has emerged from recent work is that division PG hydrolases are usually autoinhibited (1–4) and require interactions with regulatory proteins, such as EnvC and NlpD in *Escherichia coli*, to activate synchronized cleavage of the PG cell-wall. Lytic activity is thus regulated, spatially and temporally, by coupling their activation to the assembly of the cytokinetic ring (5) through the ATP-binding cassette (ABC) transporter family complex FtsEX. This membrane complex is widely conserved across diverse bacterial genera and functions as a heterodimer with FtsE in the cytoplasm and FtsX in the inner membrane (6). In *E. coli*, FtsE is one of the earliest proteins that assembles along with FtsX at the mid-cell site during cell division directly participating in this critical process by promoting septal-ring assembly in low osmolarity media (7). Indeed, FtsEX acts on FtsA to promote divisome assembly and continual ATP hydrolysis by FtsE is needed for the divisome to synthesize septal PG (8). It is worth mentioning that binding of FtsE to FtsX is independent of the nucleotide (ATP or ADP) as FtsE ATP-binding or hydrolysis mutants still localize to septal rings *in vivo* when produced together with FtsX (9). In addition, FtsEX directly recruits periplasmic EnvC to the septum via a periplasmic loop in FtsX. Interestingly, ATPase defective FtsEX variants still recruit EnvC to the septum but fail to promote cell separation in *E. coli* (10). In the divisome of G(-) bacteria, FtsE also interacts with FtsZ (11) linking the Z ring to the transmembrane protein FtsX, and it has been suggested that FtsEX may function as a membrane anchor for FtsZ utilizing ATP binding and hydrolysis to regulate Z-ring constriction (7, 11). Very recently, it has been shown in *E. coli* that recruitment of FtsE and FtsX is codependent suggesting that the FtsEX complex is recruited through FtsE interacting with the conserved tail of FtsZ (12).

The general organization of FtsEX complexes resembles an ABC transporter (7, 9, 13). This family of proteins is widespread in all forms of life and is characterized by having two copies of both the nucleotide-binding domain (NBD) and the transmembrane domain (TMD). The NBD forms dimers, which can adopt open or closed conformations. NBD are coupled to the TMD through the so-called **“**coupling helix**”** at the NBD-TMD interface [reviewed in (14)]. ATP hydrolysis in the NBD drives conformational changes that are transmitted through the coupling helices in the TMD, resulting in alternating access from inside and outside of the cell for unidirectional transport of a variety of compounds across the lipid bilayer (15). These proteins play a causative role in a number of human diseases, notably cystic fibrosis (16) and multidrug resistance by tumor cells and microbial pathogens (17). However, although FtsEX is considered a member of the Type VII family of the ABC-superfamily (18), there is no evidence that FtsEX acts as a transporter. According to this, members of this family are not typical transporters but couple ATP hydrolysis in the cytoplasm with transmembrane conformational changes to perform mechanical work in the periplasm. Thus, the FtsEX is believed to transduce a conformational signal that would couple cell division to activation of bound PG hydrolases, which vary in different bacterial species (1, 10, 19, 20). In *Streptococcus pneumoniae* FtsX interacts with the PG hydrolase PcsB (20–24) and ATPase activity by FtsE would trigger activation of PcsB (Fig. 1). PcsB is a modular protein containing a catalytic CHAP (Cysteine, Histidine-dependent Amidohydrolases/Peptidases) domain and a V-shaped coiled-coil domain (CC) (Fig. 1). Crystal structure of full-length PcsB revealed that it adopts a dimeric state in which the V-shaped CC domain of each monomer acts as a pair of molecular tweezers that clamps the catalytic CHAP domain of each partner in an inactive configuration (21). Solution studies pointed that inactive conformation is also present at low concentrations, in which monomers of PcsB would present the CHAP domain inserted in its own CC domain (21).

**Figure 1.**
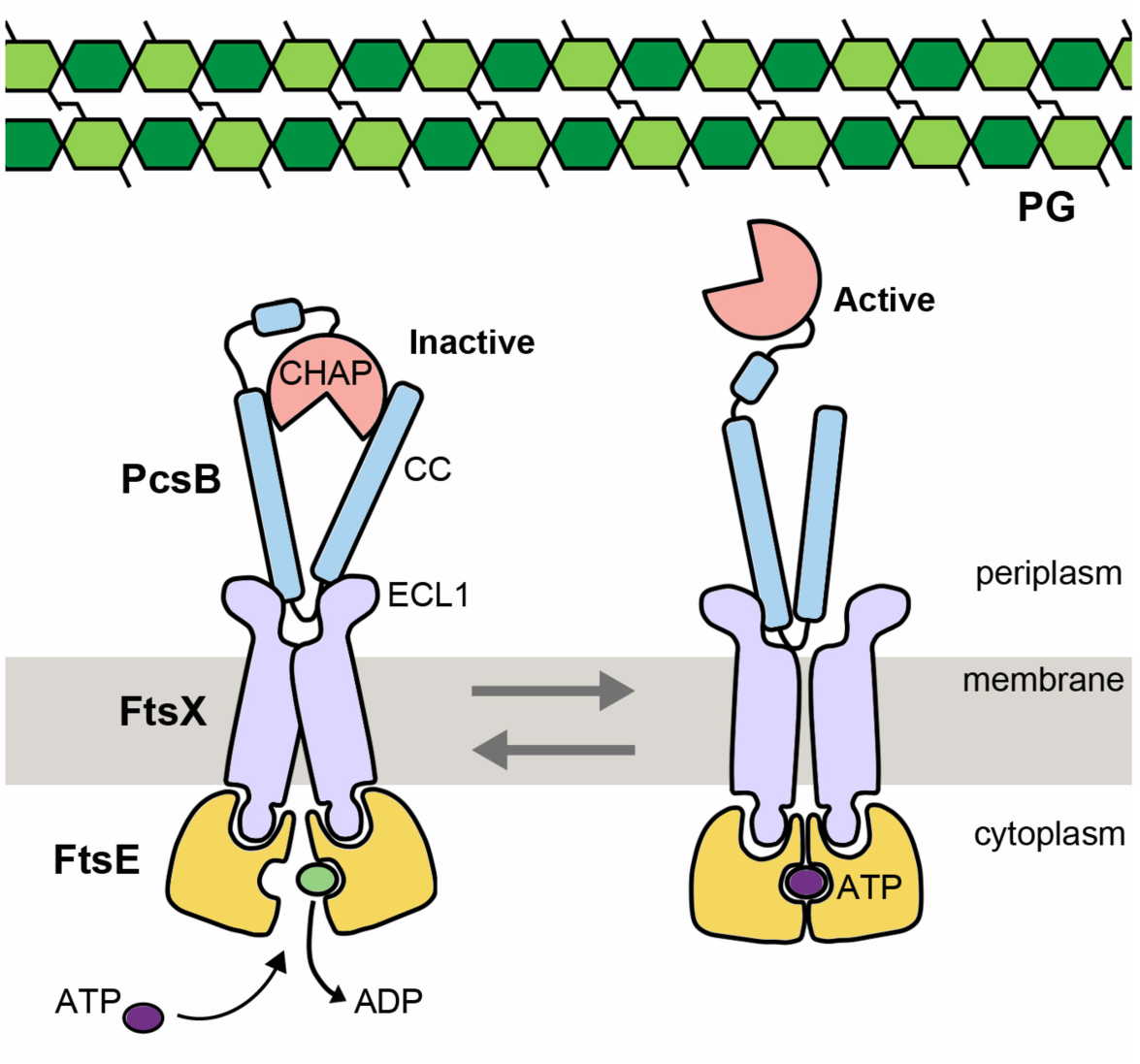
Scheme showing regulation of PG hydrolase PcsB by the FtsEX system in *S. pneumoniae*. PcsB binds to the large extracellular loop (ECL1) of FtsX independent of ATP. This binding occurs through its coiled coil (CC) domain. The stoichiometry between FtsX and PcsB is not known, here is represented as 2:1. The catalytic domain of PcsB (CHAP) is autoinhibited and activated by ATP hydrolysis of FtsEX. ATP binding leads to conformational changes in FtsEX which impact CC leading to activation of CHAP.

FtsE, FtsX, and PcsB are essential for growth in some serotype strains of *S. pneumoniae* (24–26), and their absence causes severely diminished growth and cell morphology defects in others (27, 28). In strains where FtsEX:PcsB is essential, amino acid changes that inactivate FtsE are not tolerated (24). In *E. coli, Neisseria gonorrhoeae, Aeromonas hydrophila*, and *Flavobacterium johnsoniae, ftsE* and/or *ftsX* mutants exhibit division defects, indicating that FtsEX function in cell division is conserved in these organisms (29–32). In contrast, FtsEX of *B. subtilis* has no obvious role in cell division but instead regulates entry into sporulation (33). FtsEX has been recently reported to be involved in other biological processes such as control of both the early and late stages of cytokinesis in the α-proteobacterium *Caulobacter crescentus* (34) or control of the activity of autolysin CwlO to regulate biofilm formation in *Bacillus velezensis* strain SQR9, a plant-beneficial rhizobacterium (35).

Despite the relevant biological roles exerted by FtsEX, no structural precedent for any FtsE is available to date. Here, we provide the first three-dimensional structure of FtsE (from *S. pneumoniae*) and we show that it presents all the conserved structural motifs among the NBD members of the ABC transporters family. We have obtained high-resolution FtsE structures from two different crystal forms revealing an unexpected high conformational plasticity. The results are discussed in the context of the complex that FtsE forms with the transmembrane protein FtsX. In addition, we have identified the FtsX region recognized by FtsE in the pneumococcal complex. This finding was further validated *in vivo* allowing us to propose a structural model accounting for the interaction between the two proteins.

## RESULTS

### Three-dimensional structure of FtsE from *Streptococcus pneumoniae*

FtsE was purified to homogeneity and concentrated up to 8 mg/ml (Fig. S1A) in a solution containing 25 mM Tris-HCl pH 7.5, 2 mM DTT, and 100 mM NaCl. Two different crystal forms were obtained in the same crystallization condition belonging to two different space groups, P 1 and P 2_1_ (see Materials and Methods). P 1 crystals contain one molecule per asymmetric unit and diffract up to 1.36Å resolution (Fig. S1C) while P 2_1_ crystals contain three molecules per asymmetric unit and diffract up to 1.57Å resolution (Table 1). FtsE structure was solved by *de novo* phasing with Arcimboldo (64) (see Materials and Methods). In both P 1 and P 2_1_ cases, the resulting structures correspond to monomeric FtsE and provided excellent electron density for all 230 amino acid residues of the protein (Fig. S1D). The FtsE structure has overall dimensions of ∼49.6 × 41.5 × 27.0 Å comprising two thick lobes (lobe I and II) grouped together resembling an ‘L’ shape with convex and concave sides (Fig. 2A), as previously reported for other ATPases (36). Unless otherwise indicated, the description of the monomer structure provided herein is for the highest-resolution crystal form belonging to space group P 1. In FtsE the ATP-binding pocket is located close to the end of lobe I, a subdomain which is present in many other ATPases (37). FtsE lobe I consists of a combination of α + β structures (α1 and α2 and β-sheet I, including strands β1-β2, β4-β5,) mainly formed by the packing of α1 against the antiparallel β-sheet I. The lobe II is composed only by α helices (α3−α8). Lobe I and lobe II are connected by the central β-sheet II that comprises strands β3, β6−β7−β8−β9 and β10 (Fig. 2A).

**Table 1.**
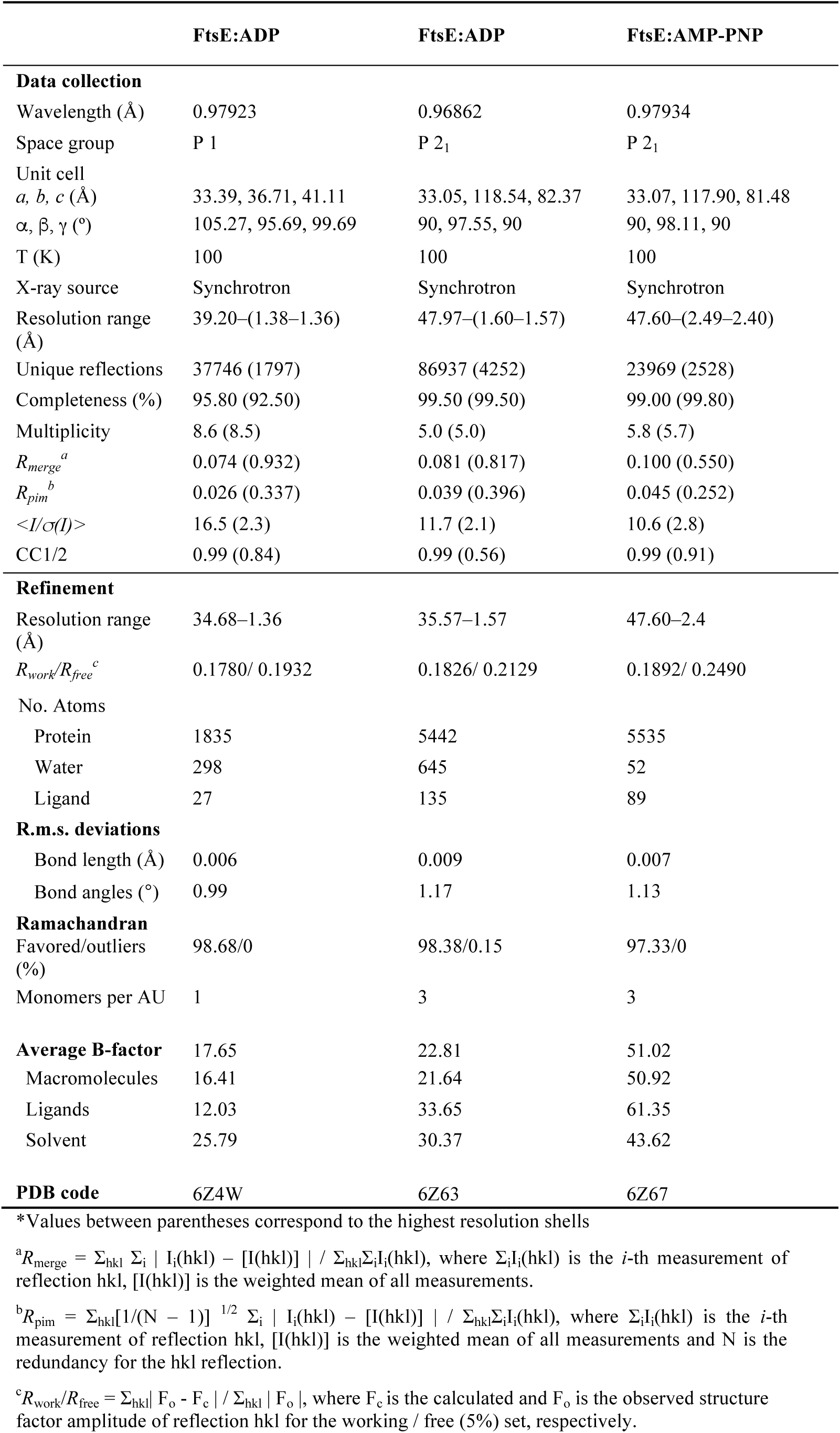
Crystallographic data collection and refinement statistics*

**Figure 2.**
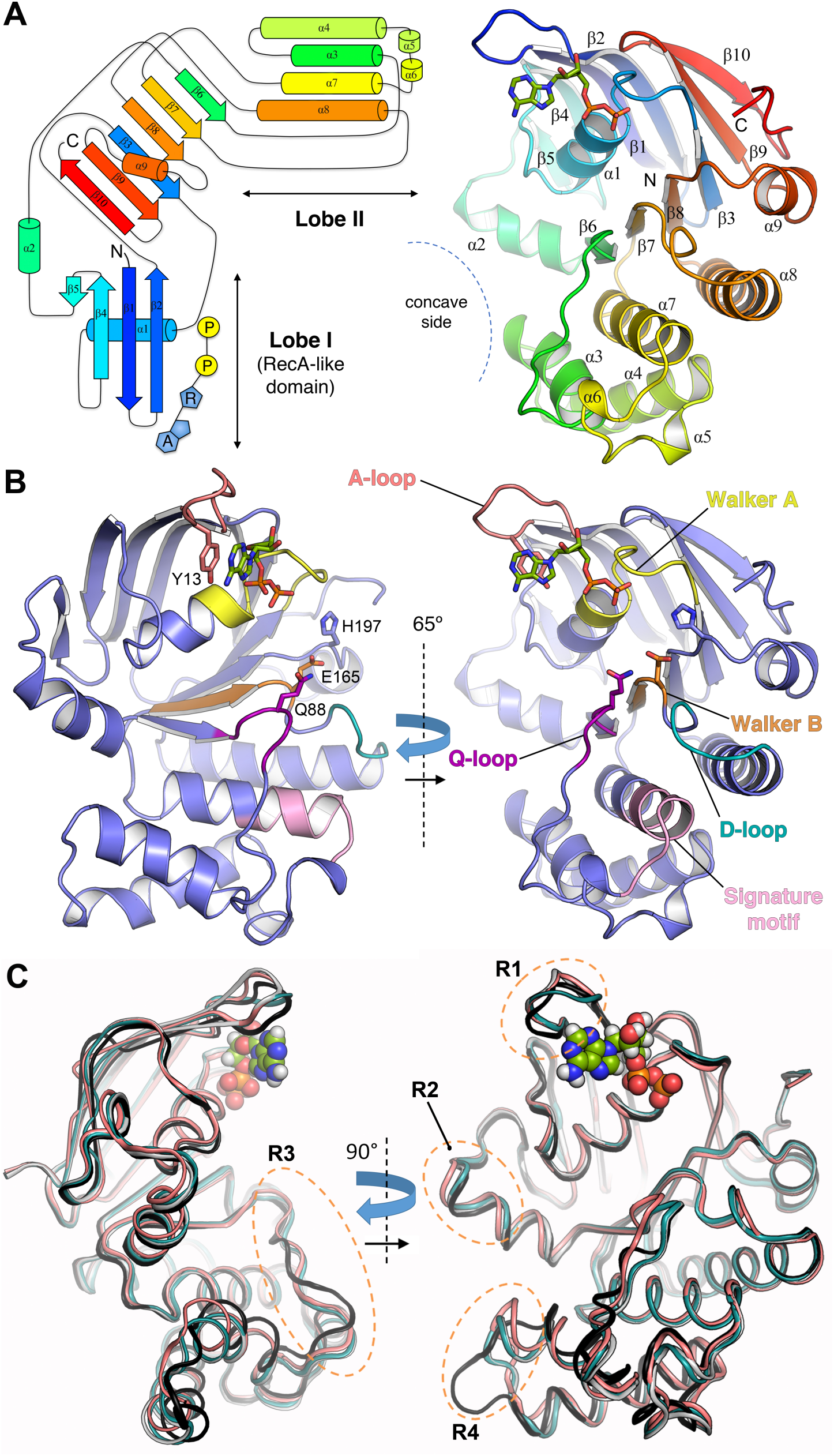
Three-dimensional structure of FtsE in complex with ADP and structural plasticity in FtsE monomer. (**A**) Left panel; topological diagram of the secondary structure of FtsE following the rainbow color code. Cylinders and arrows represent α-helices and β-strands, respectively. ADP is shown in blue and yellow (A, adenine; R, ribose; P, phosphate group). Right panel; overall ribbon representation of FtsE monomer structure. Secondary structure elements are indicated and numbered, following the same color code as indicated in A. N, N-terminus; C, C-terminus. The ADP molecule is depicted in capped sticks. (**B**) Locations of the conserved and functionally critical motifs present in the FtsE structure including Walker A/B, A-loop, D-loop, Q-loop and signature motif (colored in yellow/orange, red, cyan, magenta and pink, respectively). ADP, the catalytic glutamate (E165), the conserved glutamine (Q88) from the Q-loop and switch histidine (H197) are depicted in capped sticks. FtsE cartoon structure is displayed in two orientations at 65° of each other. (**C**) Structural superposition of the four independent FtsE monomers in complex with ADP (depicted in spheres). The protein structures are displayed in cartoon oval at two orientations at 90° of each other. Monomer P 1 is colored in red salmon, M1 is colored in black, M2 is colored in cyan and M3 is colored in pale blue. The four regions with main structural variability are indicated with dashed orange ellipsoids and labeled (R1, R2, R3 and R4, see text for details).

The NBDs present all functionally critical features belonging to the ABC transporter family including: (*i*) the Walker A motif (also known as ‘phosphate-binding loop or ‘P-loop’), which binds the α- and β-phosphates of the nucleotide, corresponds to residues 37-45 in FtsE (Fig. 2B); (*ii*) the Walker B motif is also present (residues 160-165 in FtsE) and contains the catalytic glutamate (E165), (*iii*) the ‘A-loop’ corresponds to residues 13-17 in FtsE and interacts with and positions both the adenine and ribose moieties of the nucleotide mainly by an aromatic side chain (Y13 in FtsE); (*iv*) the ‘switch histidine’ (H197 in FtsE), which stabilizes the transition-state geometry during ATP hydrolysis; (*v*) the ‘Q-loop’ (formed by residues 86-89 in FtsE), so named for a conserved glutamine (Q88 in FtsE), which provides contacts to the TMD; (*vi*) the dimerization or ‘D-loop’ (residues 168-171 in FtsE), which is involved in dimerization and has a role in coupling hydrolysis to transport [reviewed in (38)] and finally (*vii*) the α-helical subdomain that contains the “LSGGQ” signature motif (LSGGE in FtsE, including residues 140-148) which pins and orients ATP during hydrolysis and is the hallmark of the ABC transporter superfamily. In summary, FtsE presents all the features and functionally critical regions observed in NBDs of the ABC transporters.

### Structural plasticity in pneumococcal FtsE

Structural comparison of the four independent FtsE structures reported here (one monomer from the P 1 crystal form and three others, M1-M3, from the P 2_1_ crystal form) reveals significant differences among them (Fig. S2). These differences are mostly clustered in four regions (Fig. 2C). The first one (R1) is located at the A-loop (affecting residues 12-18). The second one (R2) concerns the N-terminal part of α2 (residues 71-79) that is slightly displaced among the different structures. The third region (R3, residues 86-96) presents a substantial conformational heterogeneity and affects the Q-loop, which connects β6 with α1 and is involved in the interaction with the TMD. In this region, we observe that the side chain of Y90 rotates up to ∼180° depending on the monomer (Fig. S2); as detailed below, plasticity in this region is likely related with accommodation of the coupling helix of the TMD. The fourth region (R4, residues 101-115) is affecting the C-terminal part of α3 that is even partially unfolded in monomer 1 resulting in a longer coil loop that protrudes from the monomer structure. Morphing the transition between the two most structurally different FtsE monomers (monomer from P 1 and monomer 1 from P 2_1_) illustrate the kind of movements observed between lobes I and II of FtsE (Supplementary Movie 1). The distance and the angle between the two lobes fluctuate up to ∼4 Å and 9° (considering N157 as the hinge between the two lobes), respectively, pointing out an unexpected intramolecular breathing motion within the FtsE monomer.

According to the Pfam database (39), FtsE (25795 kDa) belongs to the ABC transporters family PF0005. A 3D structural similarity search performed using DALI server revealed that the closest structural FtsE homologue is the lipoprotein-releasing system ATP-binding protein LolD from *Aquifex aeolicus* VF5 (PDB 2PCL, 39% of amino-acid sequence identity and a *rmsd* of 1.5 Å for 223 Cα atoms on superposition). Comparison with HisP, the periplasmic histidine permease of *Salmonella typhimurium* [PDB 1B0U (36)] and a well-characterized ABC transporter for this superfamily, indicates 31.4% of amino-acid sequence identity (Fig. S3A) and a *rmsd* of 2.29 Å for 224 Cα atoms. Main structural differences in FtsE compared with these structural homologues (Fig. S4) are concentrated in the ATP-binding loop (A-loop, residues 13-18) and the Q-loop (residues 86-96) both regions exhibiting structural plasticity in FtsE (regions R1 and R3 in Fig. 2); the loop 63-72 connecting β5 with α2, that in the case of HisP presents an additional β-hairpin insertion; and finally the C-terminal region that in HisP contains a helix-turn-helix motif (HisP residues 236-259), but is absent in FtsE. In addition, the C-terminus of FtsE and HisP are oriented towards the interphase of the two monomer lobes whereas in LolD is oriented in the opposite direction.

### Nucleotide recognition by FtsE

FtsE was crystallized with ADP in two different crystal forms (P 1 and P 2_1_) and with AMP-PNP, a non-hydrolysable analogue of ATP, in the space group P 2_1_ (Table 1). The nucleotide-binding site is located in lobe I near the interface between the two lobes, and presents the nucleotide sandwiched between the Walker A and B motifs (Fig. 2B). FtsE stabilizes ADP by both van der Waals and polar interactions (Fig. 3A). The Walker A (P-loop) wraps over the β-phosphate of ADP by formation of an extensive hydrogen-bond network with main-chain nitrogen atoms of G40 to K43. The position of the K43 side-chain, essential for nucleotide binding through interaction with the β-phosphate, is kept through H-bonds with main-chain oxygen atoms of G37 and P38 (Fig. 3A). The switch histidine (H197) that stabilizes the transition-state geometry in the ADP-bound state interacts with the β-phosphate through two interspersed water molecules. The Walker B motif presents the catalytic glutamate (E65) and Q88, both essential for γ-phosphate hydrolysis in HisP (36), are highly conserved in FtsE proteins from different bacterial species (Fig. S3B). Another conserved residue at the Walker B motif is D164 (Fig. S3B) that in other NBDs solved in complex with ATP, such as HisP (36), interacts with the γ-phosphate through a water molecule that occupies the position of the magnesium ion in the crystal structures of Ras-GMPPNP-Mg (40) and of the F_1_α,β-AMPPNP-Mg complex (41). This Asp residue has been suggested to interact with the divalent cation during ATP hydrolysis, by analogy with the process described for GTP hydrolysis in GTPases. The A-loop provides an aromatic side chain (Y13) that packs against the purine ring of adenine involving a π-stacking interaction and positions the base and ribose of the nucleotide. Such interaction involves an additional stabilizing force towards ADP by way of an extensive hydrophobic interaction that buries 48.43 Å^2^ of the surface area (calculated with PDBe PISA v1.52) between its aromatic ring and the adenine base of ADP. It is worth mentioning that, in the FtsE monomer, ADP molecules are further stabilized through interactions by a few residues from other monomers in the crystal. Besides, two extra ADP molecules were also found in the P 2_1_ crystal form. They increase crystal contacts (Fig. S5) but are not located in sites with physiological relevance in other ATPases.

**Figure 3.**
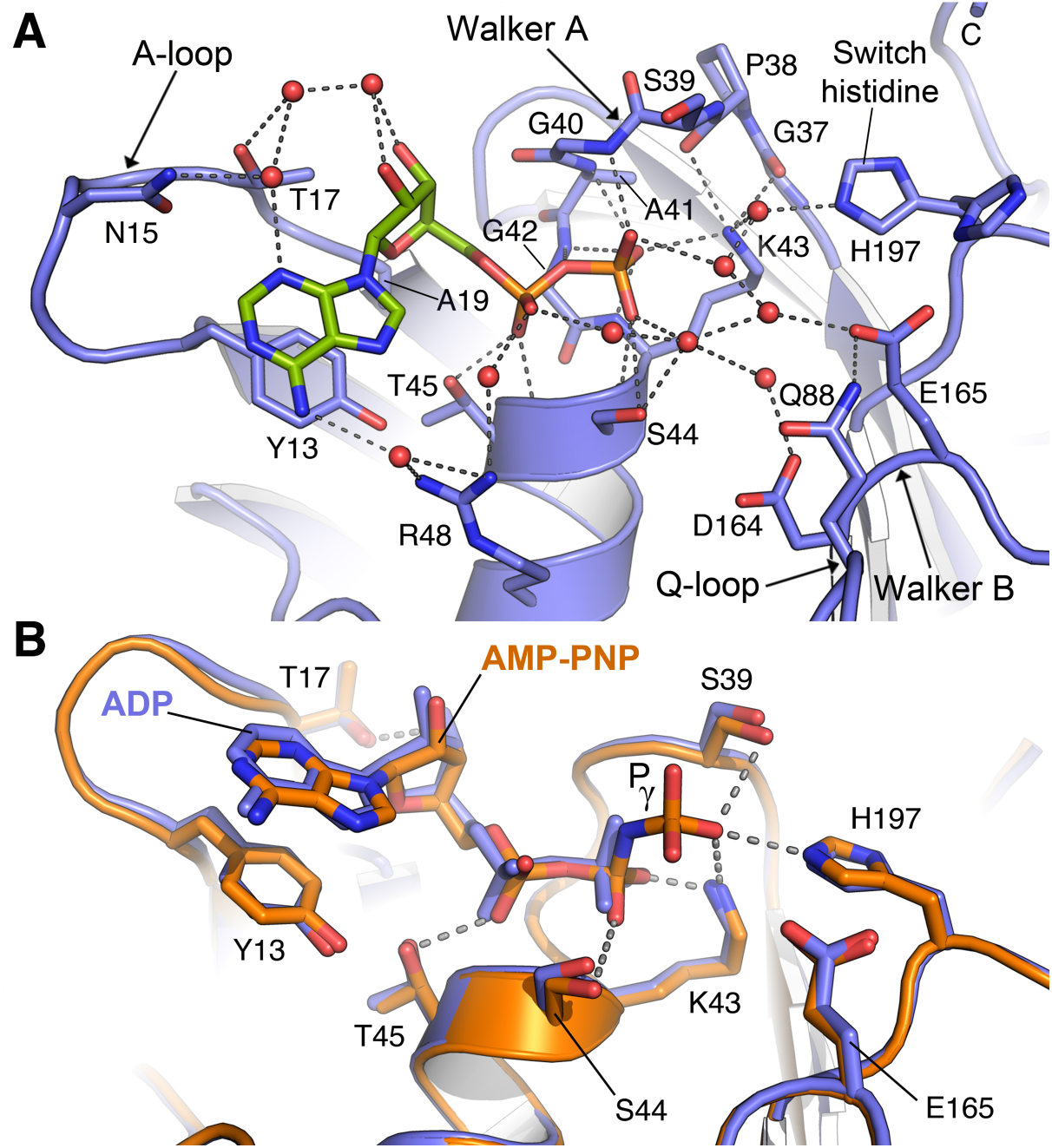
Nucleotide stabilization by FtsE. (**A**) ADP recognition at the nucleotide-binding pocket of FtsE (P1 monomer). Relevant residues involved in ADP interaction are labeled and shown as capped sticks. Water molecules are represented as red spheres. Dashed lines indicate polar interactions. Atomic interactions were determined using LigPlot (42) with default parameters. (**B**) Conformational differences between ADP-bound (colored in blue) and AMP-PNP-bound (colored in orange) FtsE monomers (M2 monomers). Residues involved in nucleotide binding, ADP (blue) and AMP-PNP (orange) are depicted in caped stick. Phosphate γ is labeled. Dashed lines indicate polar interactions.

The FtsE:AMP-PNP complex was obtained by soaking with the P 2_1_ FtsE:ADP crystals (Table 1). The ATP analog was found in two out of the three independent monomers of the P 2_1_ crystal. Structural comparison between our ADP and AMP-PNP complexes revealed that there were no relevant structural rearrangements directly associated to AMP-PNP binding, *e*.*g*. monomer M2 in complex with ADP or with AMP-PNP were nearly identical (*rmsd* of 0.22 Å for 223 Cα atoms) (Fig. 3B). In the presence of AMP-PNP, K43 interacts with both β-phosphate and γ-phosphate. The γ-phosphate is further stabilized by a direct interaction with the switch histidine (H197) and with S39 (Fig. 3B). The catalytic glutamate (E165) is positioned directly adjacent to the γ-phosphate consistent with a role in ATP hydrolysis. In conclusion, the relevant residues to recognize the ATP molecule are properly oriented and no relevant changes were required for nucleotide exchange.

### Model for FtsE dimerization

FtsE performs its catalytic activity *in vivo* as a dimer. The availability of structures of ABC transporters in the dimeric state allowed us to build a structural model accounting for the global architecture of the FtsE dimer in the two main, ADP and ATP, physiological states. For the ADP state we used our FtsE:ADP complex and the crystal structure of tripartite-type ABC transporter MacB from *Acinetobacter baumannii* in the ADP state [PDB 5GKO (43)] as template (Fig. S6A). Superposition of the NBD of MacB with FtsE:ADP complex results in an excellent fitting with a *rmsd* value of 1.0 Å for 174 Cα atoms (Fig. S6B). In this model, FtsE dimerizes in a “head-to-tail” orientation, where each lobe II mutually binds to lobe I of the opposing monomer (Fig. 4A). The interactions between the two subunits in the obtained homodimer model are extensive, burying about 11493 Å^2^ (calculated with PDBe PISA) of accessible surface per subunit. In the FtsE dimer, the ADP nucleotide is stabilized only by a monomer and thus there is no interaction between the nucleotide and the dimeric partner, in agreement with previous observations for ABC transporters (Fig. 4A).

**Figure 4.**
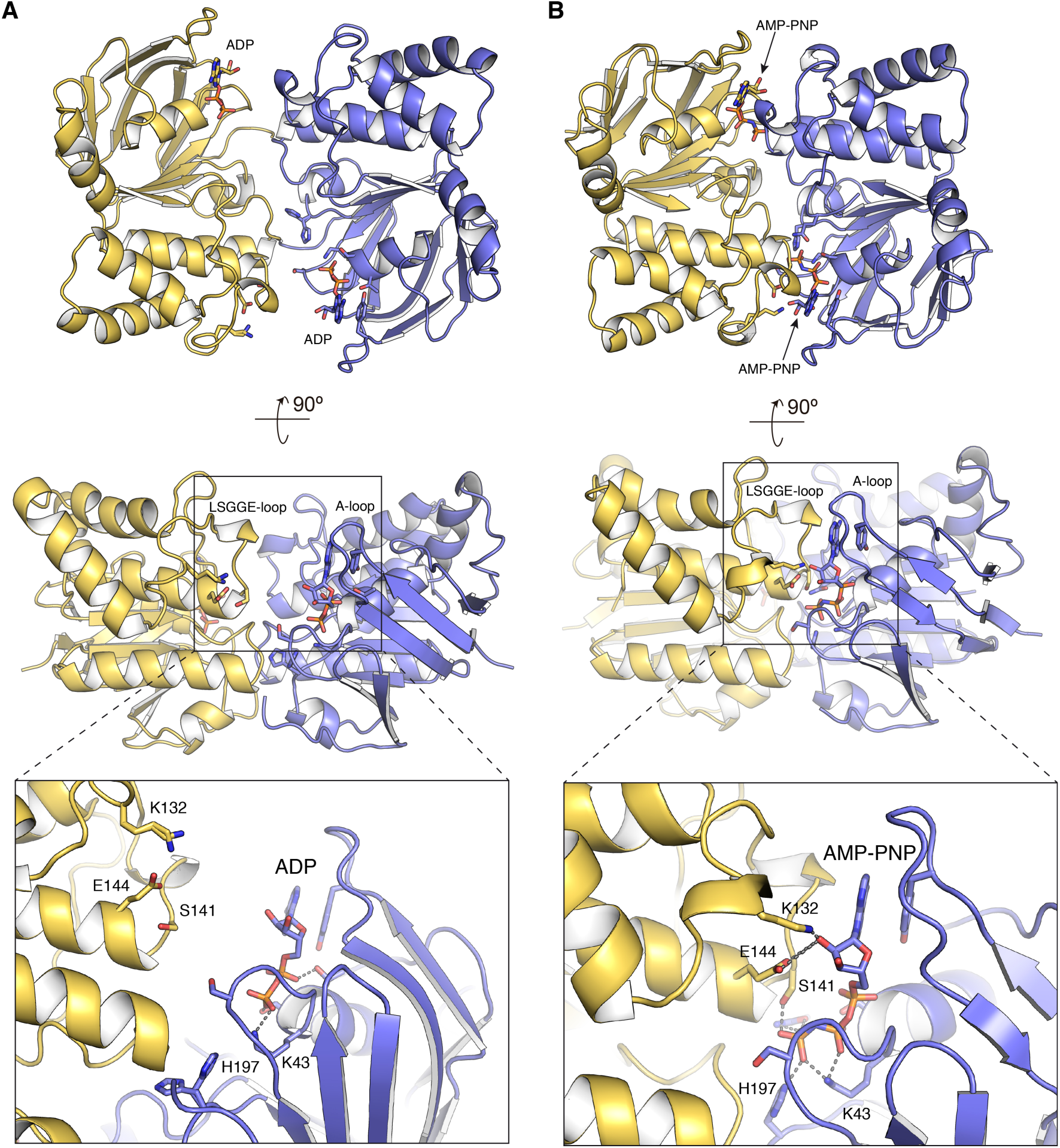
Model for FtsE dimerization in the ADP and ATP states. (**A**) FtsE dimer in the ADP-bound conformation. The FtsE:ADP dimer (monomers colored in blue and yellow) as obtained by structural superposition with the protein MacB from *A. baumannii* [PDB 5GKO (43)]. Boxed region at the bottom shows a detailed view of the nucleotide-binding site in the dimer. **(B)** FtsE dimer in the ATP-bound conformation. The FtsE:AMP-PNP dimer (monomers colored in blue and yellow) as obtained by structural superposition of FtsE:AMP-PNP with the protein MacB:ATP from *A. actinomycetemcomitans* [PDB 5LJ6 (44)]. Boxed region at the bottom shows a detailed view of the nucleotide-binding site in the ATP dimer. ADP and AMP-PNP molecules and side-chains of relevant residues are represented as capped sticks and labeled. Dashed lines indicate polar interactions of the nucleotide with protein residues.

To create a model for the FtsE dimer in the ATP state we used the FtsE:AMP-PNP complex and the crystal structure of the *Aggregatibacter actinomycetemcomitans* MacB bound to ATP [PDB 5LJ6 (44)] as template (Fig. S6C). While direct superposition of both structures reveals an overall nice fitting (*rmsd* value of 2.5 Å for 212 Cα atoms), we observed that MacB:ATP structure presented a more compact structure for the nucleotide-binding domain (both lobes closer to the central β-sheet II) than in FtsE. We then prepared a composite model in which FtsE was fragmented in three subdomains: lobe I-core (residues 1-86, 159-164 and 191-228), lobe II (residues 87-158) and α-helix 8 (residues 165-190). Superposition of each region onto the MacB:ATP structure presents excellent *rmsd* values (0.8 Å for 125 Cα atoms in lobe I-core, 0.78 Å for 64 Cα atoms in lobe II and 0.5 Å for 26 Cα atoms in α-helix 8) (Fig. S6C and S6D) indicating rigid-body movements of these regions around the central β-sheet in the ATP state. In the ATP conformation there is a closer contact between FtsE monomers with the LSGGE-loop of the dimeric partner sandwiching the AMP-PNP bound to the A-loop of the monomer (Fig. 4B). In this model, FtsE residues K132 and E144 of the partner would interact with the ribose moiety (Fig. 4B); equivalent residues in MacB (K136 and Q148) also interact with ribose in the ATP complex. It is worth mentioning that in MacB the γ-phosphate is further stabilized by H-bonds though main chain atoms of G147, A173 of the partner (G142, N169 in FtsE); besides, our model predicts potential implication of the side-chain of S141 in stabilization of the γ-phosphate (Fig. 4B).

In addition, residues involved in direct protein-protein interactions across the dimer interphase previously described for other NBDs in the dimeric state such as MalK [PDB 1Q12 (45)], are conserved in pneumococcal FtsE. These residues involve S38 (S39 in FtsE) that forms strong hydrogen bonds with R141 and D165 (R147 and D171 in FtsE) and H192 (H197 in FtsE) that makes hydrogen bond and van der Waals interactions with N163, L164 and D165 (N169, L170 and D171 in FtsE).

### Identification of the FtsX region interacting with FtsE in *S. pneumoniae*

FtsX is an integral membrane protein with cytoplasmic amino and carboxyl termini, four transmembrane segments, and one large and one small <u>e</u>xtra<u>c</u>ellular loop (ECL1 and ECL2, respectively) (Fig. S7A). We recently reported the structural and functional characterization of the large extracellular domain ECL1 from *S. pneumoniae* (46) connecting the first two transmembrane helices (Fig. 5A and Fig S7A). In the periplasmic space the ECL1 interacts with PcsB protein, which harbors a CHAP domain with the endopeptidase activity required for normal cell division. In the cytoplasm FtsX interacts with FtsE, but the FtsX regions involved in the formation of the complex in *S. pneumoniae* are still not known. As described before, FtsE presents a high sequence homology with the nucleotide-binding domains of ABC transporters, and also between FtsE proteins from different bacterial species (Fig. S3B and S3C). Conversely, sequence conservation among FtsX proteins from different bacterial species is rather low (Fig. S7B). This feature probably reflects the fact that FtsX binds PG hydrolases, which vary among different bacterial species (1, 10, 19, 20). Surprisingly, we identified a region of 22 residues with the sequence-NTIRITI<u>ISRSREIQIMRLVGA</u>- (residues 201-222 in *S. pneumoniae* strain R6) predicted as cytosolic and presenting a significant degree of conservation among the different FtsX sequences that we analyzed (Fig. S7B). This region is located between TM2 and TM3 and therefore would likely correspond to the unique cytoplasmic loop present in FtsX. We speculated that the sequence conservation of this region could be due to the interaction with its cytoplasmic partner FtsE. To test this hypothesis, we labeled FtsE with a fluorescent dye and performed Microscale thermophoresis (MST) analysis (see Material and Methods) that show that peptide ISREIQIMRLVGA (which sequence is underlined in the 22 residues long sequence showed above) interacts with FtsE with low affinity (Kd of ∼82 nM, Fig. S7C). All our attempts to co-crystallize FtsE with this FtsX fragment failed, but a three-dimensional model for this sequence was generated with the Swiss-Model server (47) that identified PDB 5NIK as the best template sharing 32.26% of sequence identity with the query. This structure, nicely corresponds to the MacB component of the MacAB-TolC ABC-type tripartite multidrug efflux pump (48), and the region of this structure identified as template corresponds to an N-terminal helix of roughly 20 residues that precedes TM1, and skirts along the inner leaflet of the cytoplasmic membrane before making an abrupt turn at nearly a right angle into the interior of the lipid bilayer. This N-terminal helix corresponds to the ‘coupling helix’ found in other ABC transporters (49). A groove in the FtsE surface forms the contact interface with the coupling helix of the transmembrane domain (TMD). Although the coupling helix is not the only contact between TMDs and NBDs, it is the only architecturally conserved contact among distinct TMD folds and provides the majority of contacts between domains. Based on this three-dimensional model for the FtsX cytoplasmic loop, we identified some potential residues promising for a mutagenesis study in order to demonstrate that indeed this FtsX region is interacting with FtsE in *S. pneumoniae*. The selected residues were E213, I216, L219 and V220. Residues I216, L219 and V220 are hydrophobic that would be buried in a cavity present in the FtsE structure that accommodates the FtsX ‘coupling helix’ (Fig. 5). E213 would be protruding outwards from the ‘coupling helix’ establishing polar interactions with residues from FtsE (*i*.*e* K95, Fig. 5B) and far away from the FtsE cavity.

**Figure 5.**
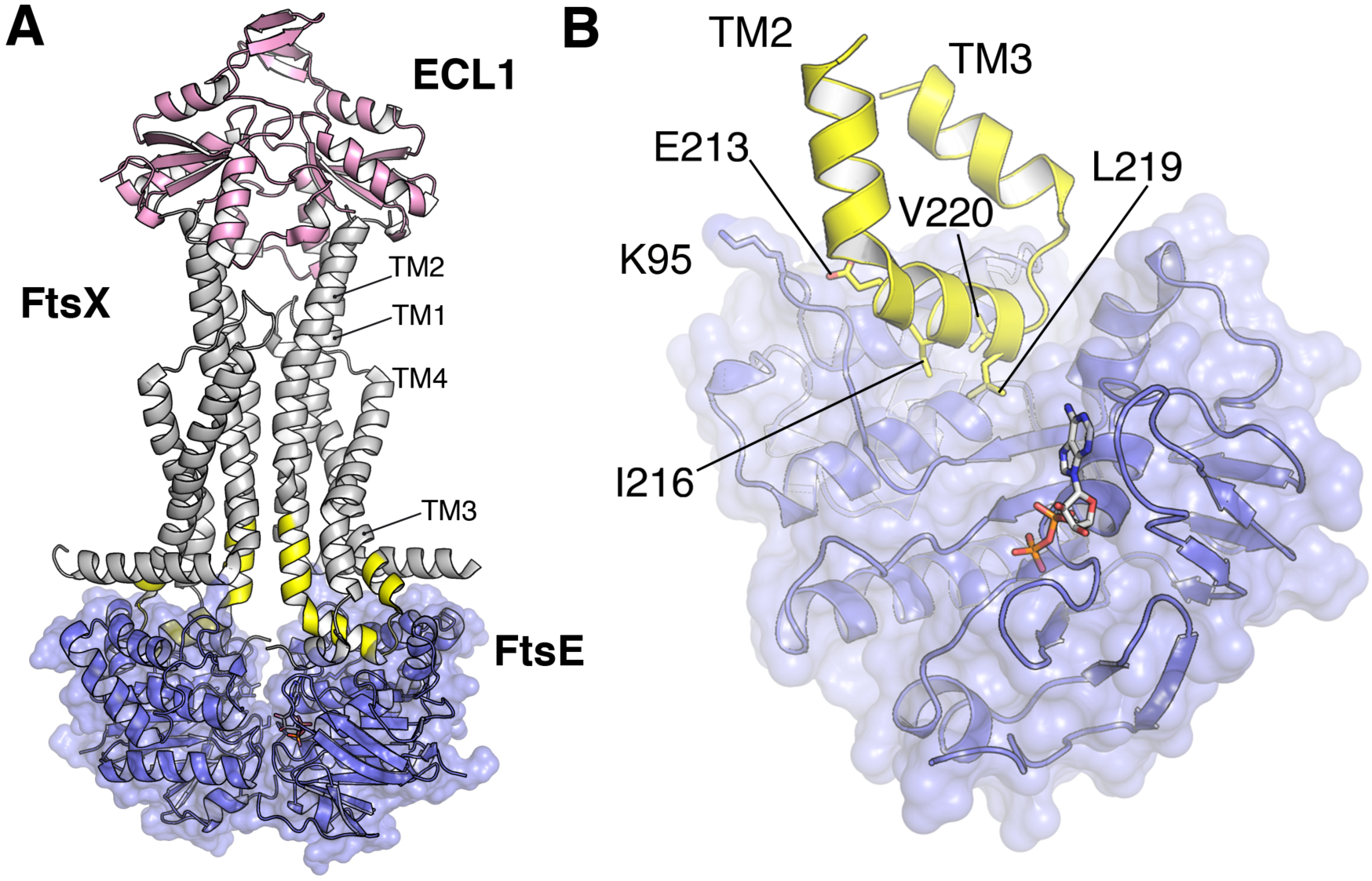
Identification of the FtsX region interacting with FtsE. (**A**) Full-model for FtsEX complex with the structure of FtsE (this work) colored in blue, the crystal structure of the large extracellular domain of FtsX (ECL1, colored in pink) from *S. pneumoniae* [PDB code 6HFX (46)] and the transmembrane domain (colored in grey) as observed in MacB from *A. Baumannii* [PDB code 5GKO (43)]. ECL1 from pneumococcal FtsX was structurally superimposed onto the ECD of MacB. (**B**) Three-dimensional model of the FtsX putative coupling helix (yellow cartoon) obtained with The Swiss-Model server (47) using PDB 5NIK (48) as a template. Relevant residues selected for mutagenesis have been labeled and depicted in sticks. FtsE K95, a residue that could interact with FtsX E213, is shown in sticks. FtsE monomer is shown in blue cartoon with semitransparent surface. ADP is depicted in sticks.

### FtsE interacts with the cytoplasmic region between TM2 TM3 of FtsX in *S. pneumoniae* and this interaction is essential for cell growth and proper morphology

Once the potential interacting region between FtsX and FtsE was identified, we next determined the degree to which this interaction interface contributes to pneumococcal viability. For this, we decided to target the above-mentioned residues for substitution with lysine. Construction of the indicated mutants was done by using overlap extension PCR and the resulting sequences were inserted at the native locus in the *S. pneumoniae* genome by double cross-over via homologous recombination after transformation (see Materials and Methods). Given the essentiality of the interaction of FtsE and FtsX, mutagenesis of these residues might have a lethal effect. Thus, we used a strain containing an ectopic copy of *ftsX* under the control of the inducible P_*comR*_ promoter (see Materials and Methods) that allows the depletion of ectopic FtsX expression by using the ComRS gene expression/depletion system. Subsequently, mutants were inspected for growth or morphology defects. By growing the cells without ComS inducer in a two-fold dilution series, the level of intracellular ComS is gradually reduced through cell division and metabolism. To test the functionality of different FtsX mutants, ectopic FtsX was depleted in the following genetic backgrounds: Δ*ftsX, ftsX*_*wt*_, *ftsX*^*E213K*^, *ftsX*^*I216K*^, *ftsX*^*L219K*^ and *ftsX*^*V220K*^. All mutants tested presented phenotypic defects similar to those of cells in which *ftsX* was depleted, typically resulting in reduced growth and cell chaining (Fig. 6A and 6B). Comparison of their cell shape distributions (length/width) showed that the most severely affected mutants, I216K and V220K, also resembles the *ftsX* depleted cells in being less elongated than *wt* cells, and having a significant number of bloated cells (Fig. 6B and 6D). The L219K mutant also displayed shorter cells and bloated cells, whereas the E213K mutant did not. However, when comparing the mean cell area (µm^2^) all FtsX mutants were in general smaller than the *wt* (Fig. 6E), despite that several bloated cells were seen for the non-viable mutants. We also constructed Flag-tagged versions of the point mutated FtsX proteins to check that their expression levels and stability were similar to *wt* FtsX when expressed from the native promoter (Fig. 6C). This analysis revealed that all mutants were expressed at nearly *wt* FtsX levels indicating that the observed differences were not due to misexpression of FtsX. However, the levels of E213K appeared to be somewhat reduced compared to the others. We do not believe this is a critical issue since this mutant was viable, although it displayed chaining and a slight growth reduction, the cells length/width ratios were not significantly different from *wt* cells. The I216K and V220K mutants, which were lethal, are expressed at *wt* levels confirming that it is not the lack of FtsX protein that caused this phenotype, but the loss of interaction with FtsE. The E213K mutant has the most permissive phenotype, probably due to its location protruding outside the hydrophobic cavity present in FtsE.

**Figure 6.**
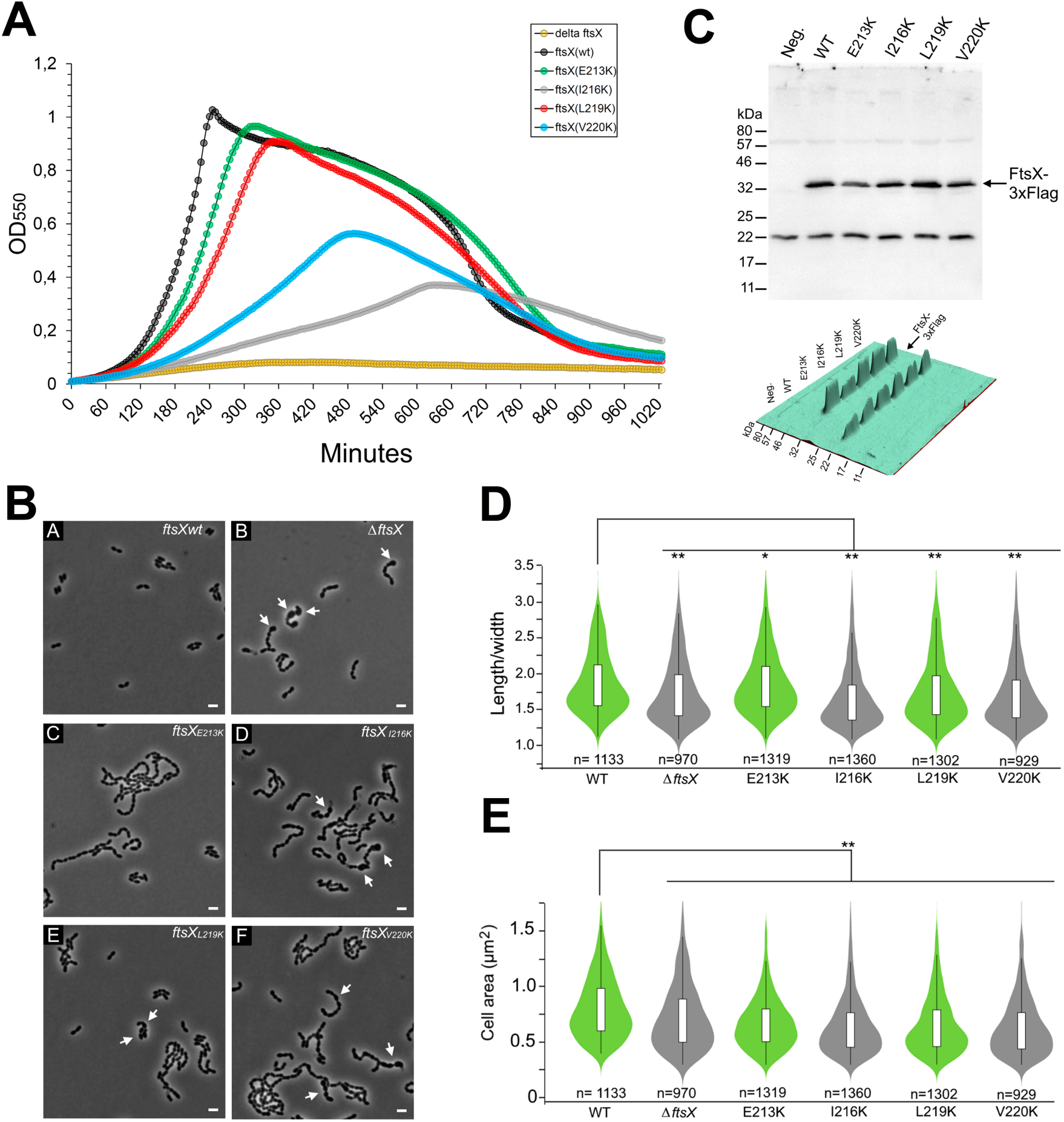
**(A)** Depletion of ectopic FtsX expression. Representative growth curves from dilution eight (128 fold diluted) is shown. (**B**) Morphology of pneumococcal strains expressing FtsX^E213K^, FtsX^I216K^, FtsX^L219K^ and FtsX^V220K^ compared to wild type and a Δ*ftsX* strain. All strains were depleted of an ectopic copy of *ftsX*, to test the function of the different *ftsX* versions placed in the native site in the genome. Arrows indicate bloated cells. Scale bars are 2 µm. (**C**) Stability of point mutated versions of FtsX. Flag-tagged versions of FtsX, FtsX^E213K^, FtsX^I216K^, FtsX^I219K^ and FtsX^V220K^ expressed from the native *ftsX*-locus were detected in whole cell extracts using immunoblotting. Expected molecular weight (MW) for FtsX is 34.2 kDa. The Flag-negative strain RH425 was used as control for background signals. A 3D-visualization of the signal intensities is shown at the bottom. (**D** and **E**) Violin plots showing the mean cell/width distribution and cell area (µm^2^) of *wt* cells and the *ftsX* mutants. (**D**) The viable mutants are shown in green and the lethal mutations in grey. Comparison of the mean length/width ratio of the *wt* (1.87 ± 0.42) with Δ*ftsX* (1.74 ± 0.44), *ftsX*^*E213K*^ (1.86±0.43), *ftsX*^*I216K*^ (1.66±0.43), *ftsX*^*L219K*^ (1.76 ± 0.45) and *ftsX*^*V220K*^ (1.69 ± 0.44) cells showed that the ratio of the Δ*ftsX, ftsX*^*I216K*^, *ftsX*^*L219K*^ and *ftsX*^*V220K*^ mutants were significantly lower than *wt* cells, with the non-viable mutants I216K and V220K showing the lowest length/width ratios. Mutant E213K did not display a significantly different ratio compared to *wt* cells. (**E**) Comparison of the mean cell area of *wt* cells (0.81 ± 0.28) with the *ftsX* mutants Δ*ftsX* (0.72 ± 0.30), *ftsX*^*E213K*^ (0.68 ± 0.24), *ftsX*^*I216K*^ (0.64 ± 0.26), *ftsX*^*L219K*^ (0.65 ± 0.26) and *ftsX*^*V220K*^ (0.64 ± 0.27). All *ftsX* mutants were significantly smaller than *wt* cells. The number of cells counted is indicated. P-values were obtained relative to *wt* using one-way Anova analysis. *P>0.05, **P<0.001.

Taken together, these results reveal that the cytoplasmic region of FtsX that links TM2 and TM3 is critical for FtsX function *in vivo* and confirm the functional importance of the physical interaction of FtsX and FtsE.

## DISCUSSION

FtsEX is member of a subclass of ABC transporters using mechano-transmission to accomplish work in the periplasm. Two ABC transporters are closely related to FtsEX having the same transmembrane topology and with an ECD located between the first and second membrane-spanning helices. These include MacB and LolCDE, both of which have four TM segments and assemble into ABC transporters as dimers with a total of eight TM segments (50–52). LolC and LolD work together with the cytoplasmic ATPase LolE to transfer lipoproteins from the outer surface of the cytoplasmic membrane to the periplasmic chaperone LolA (53, 54). MacB helps to secrete heat-stable enterotoxin II from the periplasm across the outer membrane (55). In both cases, their activity does not involve moving a substrate across a membrane, and the TM segments serve to transmit ATP-driven conformational changes rather than forming a substrate channel, a process known as mechano-transmission (44). The FtsEX complex is expected to behave in this same manner, coupling cytoplasmic ATP hydrolysis with extracellular conformational changes that drive the activation of the PG hydrolytic activity of PcsB during bacterial division (Fig. 1).

In this work we present the first three-dimensional structure of FtsE and we unveil the interactions with its membrane partner FtsX through the region connecting TM2 and TM3. We show that FtsE presents all the conserved structural features (WalkerA/B, P-loop, Q-loop, D-loop, A-loop and signature motif) associated to the NBD domains of ABC transporters superfamily.

The four independent FtsE structures, here provided, revealed an unexpected conformational plasticity that is translated into an intramolecular breathing motion within the two FtsE lobes (Supplementary movie 1). FtsE plasticity is observed in four regions; R1 that affects the tip of the A-loop responsible for recognition of the adenine ring of the nucleotide, R2 (residues 71-79), R3 (residues 86-96) that includes the Q-loop (residues 86-89) and finally the region R4 (residues 101-115).

The disposal of similar NBDs structures in complex with their corresponding TMDs (43, 44) allowed us to generate a model for the pneumococcal FtsE dimer both in the ADP- and ATP-bound conformations in complex with the FtsX coupling helix. Interestingly, three out of the four FtsE regions exhibiting conformational plasticity (regions R2, R3 and R4) shape the cavity in which the FtsX coupling helix would be located (Fig. 7A and 7B). In agreement with its dynamic behavior, these FtsE regions present above-average temperature factors that, together with their close proximity with the TMD, will favor mechano-transmission to FtsX. The bottom of the cavity is mainly composed by hydrophobic residues, among them F87 (Fig. 7A). The relevance of these hydrophobic interactions is in agreement with our observation that FtsX mutants affecting hydrophobic residues at the cytoplasmic loop (I216K, L219 and V220K) have a dramatic phenotype affecting cell growth and division.

**Figure 7.**
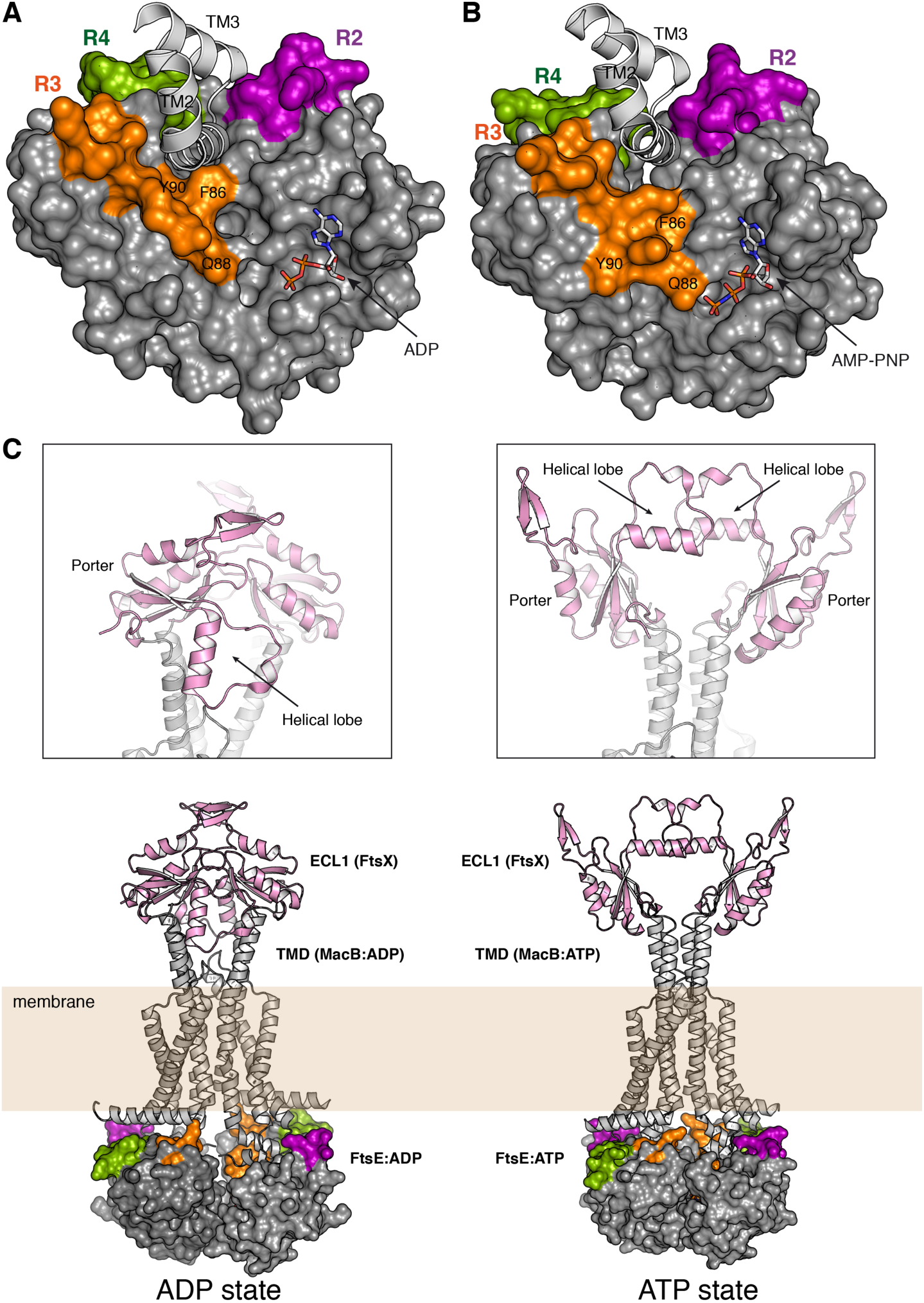
(**A**) Model of the FtsE:ADP complex with FtsX coupling helix. Molecular surface of the FtsE:ADP complex (monomer P1) colored in grey with regions R2 (residues 71-79), R3 (residues 86-96) and R4 (residues 101-115) colored in purple, orange and green respectively. TM helices from MacB from *A. baumannii* [PDB 5GKO (43)] represented as white cartoon. (**B**) Model of the FtsE:ATP complex with FtsX coupling helix. Molecular surface of the composite model of FtsE:AMP-PNP complex obtained by superimposition onto the *A. actinomycetemcomitans* MacB bound to ATP [PDB 5LJ6 (44)]. Regions R2 (residues 71-79), R3 (residues 86-96) and R4 (residues 101-115) highlighted in purple, orange and green respectively. TM helices from MacB:ATP represented as white cartoon. ADP and AMP-PNP molecules represented as capped sticks. Some relevant residues (see main text for details) are labeled. (**C**) Full-model for the pneumococcal FtsEX complex in the ADP and ATP states. The TM helices correspond to the MacB complex in ADP or ATP states (PDB code 5GKO and 5LJ6 respectively). Position of the pneumococcal ECL1 of FtsX (PDB code 6HE6) was obtained by direct superimposition of the Porter subdomain of FtsEx onto the Porter subdomain of MacB. Details of the arrangement of the Helical lobes of FtsX in ADP and ATP states are shown in the boxed areas.

Other conclusions can be extrapolated from our structures; we have observed that while the relaxed structure of the FtsE:ADP complex nicely fits the equivalent complex in MacB dimer; the monomer of the FtsE:AMP-PNP complex requires rigid-body adjustment of some regions around the central β-sheet in order to get a perfect fitting with the more compact structure of the MacB:ATP dimer. Considering the high sequence homology among the NBD domains of ABC transporters, this result indicates that protein-protein interactions in the dimer are required for full-recognition of the ATP nucleotide (as we predicted by our ATP-bound FtsE dimer, Fig. 4B), and also to reach the final 3D structure *in vivo*. Observed plasticity in FtsE would favor such protein-protein mediated changes as also occurs in our crystal structures.

Disposition and conformation of FtsE R2-R4 regions is different in ADP or ATP states (Fig. 7A and 7B) thus altering the global shape of the FtsX-binding cavity. Structural changes observed in the backbone or in the side-chain conformation of some residues (such as Y90 residue) in these regions would correlate with the changes between ADP and ATP conformations in FtsE. For instance, in our FtsE:AMP-PNP complex the γ-phosphate is far from the Q88 residue of the Q-loop (> 6Å), however our composite model for the same structure in the dimer indicates that Q88 would directly recognize the γ-phosphate (Fig. 7B) as also occurs in the structure of the MacB:ATP dimer (44). In conclusion, FtsE regions R2-R4 would play a relevant role in the mechanic transmission from the cytosol through the membrane in order to regulate lytic activity in the periplasm.

*S. pneumoniae* and most other Gram-positive bacteria synthesize a complete septal cross-wall before cell separation takes place at the end of the cell cycle. In Gram-negative bacteria, on the other hand, synthesis and splitting of the septal cross-wall and invagination of the outer membrane occur in tandem (56). Hence, the constrictive mode of cell division displayed by Gram-negative bacteria requires a tight coupling between peptidoglycan synthesis and hydrolysis during septation. In *E. coli*, FtsEX, which localizes to the Z ring, has been shown to play and important role in this process (12). In this Gram-negative species, FtsEX is involved in assembly of the divisome, activation and regulation of peptidoglycan synthesis, and recruitment and activation of cross-wall splitting hydrolases (6). Interestingly, FtsEX is not involved in cell division in *B. subtilis*. Instead, FtsEX controls the peptidoglycan hydrolase CwlO, which plays an important role in cell wall expansion during cell elongation (57). In *S. pneumoniae*, FtsEX controls the peptidoglycan hydrolase PcsB, which is essential for cross-wall splitting and cell separation in some serotypes and required for normal cell separation in others (21). Of note, synthesis and splitting of the septal cross-wall are separate in time and space in *S. pneumoniae*. The separation of daughter cells takes place at the end of the cell cycle, well after the synthesis of the septal cross-wall has been completed. Hence, it is highly unlikely that the FtsEX-PcsB complex is associated with the Z ring and the core divisome in *S. pneumoniae*. We have previously reported that pneumococci form very long chains when PBP2b, a key component of the elongasome, is depleted. Furthermore, transmission electron microscopy analysis of these cells revealed that their septal cross-wall has not been split (58). Thus, when the elongasome is no longer functional, separation of daughter cells mediated by the FtsEX-PcsB complex does not take place. This observation shows that the elongasome must be properly assembled before the FtsEX-PcsB complex becomes active, suggesting that the signal that triggers cross-wall splitting and daughter cell separation in *S. pneumoniae* is linked to elongasome assembly and the initiation of lateral peptidoglycan synthesis. To further explore which FtsX region is recognized by FtsE in *S. pneumoniae* we have performed an *in vivo* mutagenesis study. Our results clearly pinpoint the FtsX cytoplasmic loop (residues 201-222) linking TM2 and TM3 as the FtsE binding site. Our results are in agreement with a recent work in which the authors show that a mutations affecting this FtsX region in *E.coli* markedly reduced the ability of ftsEX to complement a *ΔftsEX* mutant (12).

How the mechano-transmission works for FtsEX to affect PcsB is a central question that still remains unanswered, *i*.*e*., how the ATP *vs* ADP version of FtsE at cytosol affects ECL1 at periplasm, which then affects PcsB? While the final answer will come, likely, from the 3D structure of the binary and ternary complexes (FtsEX and FtsEX:PcsB), present structural information can provide some insights on this matter. The structure of pneumococcal ECL1 of FtsX (46) presents a Helical lobe and the so-called Porter subdomain which is remarkable similar to that of *E. coli* MacB (1, 44). The Porter subdomain is present in all members of Type VII ABC superfamily and is likely an intrinsic part of the mechanotransmission apparatus (18). Structural superposition of the Porter subdomains in the pneumococcal ECL1 and in MacB, in both ATP and ADP states, has allowed us to generate a three-dimensional model accounting for the arrangement of the FtsX ECL1 between both states (Fig. 7C). Interestingly, depending on the ADP or ATP state, the Helical lobe of ECL1 changes completely its orientation. In the ATP state the Helical lobes from each ECL1 are very close and located face to face, while in the ADP state the Helical lobes of each ECL1 are in opposite directions (Fig. 7C). We previously reported that ECL1 Helical lobe mediates a physical interaction with PcsB (48). Thus, this difference between ATP and ADP states would dramatically alter the FtsEX-PcsB interaction. This conclusion could be applied to other bacteria like *E. coli*, where it was shown that this same FtsX lobe (referred to as lobe X in Gram negative bacteria (6)) is required for binding to EnvC (59). In our prediction, the secondary structural elements that make up ECL1 are not altered and it would imply just a rotation of ECL1 dictated by the scissors-like action of the transmembrane domains driven by the dimerization of the FtsE, thus providing the driving force regulating lytic activity in the pneumococcal division.

## Material and methods

### Cloning, expression and purification of protein FtsE

We fused FtsE at its N-terminus to a 6xhis-followed by a TEV protease cleavage site in vector pETDuet™-1 (Novagen) that allows tunable expression of the fusion protein from a T7 promoter induced by isopropyl-β-D-1-thiogalactopyranoside (IPTG). Briefly, the DNA fragment corresponding to *ftsE*, flanked by EcoRI and HindIII restriction sites, was amplified by PCR from genomic DNA of *S. pneumoniae* strain R6 (primers Ftse1 and Ftse2, Table S2), digested with restriction enzymes EcoRI and HindIII, and cloned into equivalent sites of vector pETDuet™-1 at the Multi Cloning Site 1. *Escherichia coli* DH5α cells were transformed with this ligation mixture, and ampicillin-resistant transformants were selected. The resulting construct was verified by Sanger sequencing and used to transform *E. coli* Lemo21. This strain (BL21DE3/pETDuet-1-*6xhis-TEV-ftsE*) was grown until OD_600_ of 0.5 and his-FtsE expression was induced with 0.3 mM IPTG (Promega) during 3 hours at 28°C. Cell pellets were resuspended in Buffer E (50 mM Tris-HCl pH 7.5, 500 mM NaCl, 20 mM imidazole, 2 mM ATP and 10% glycerol) containing 1 mmol/L PMSF and DNase I (Roche Diagnostics Corp., IN, USA) to 5 µg/mL. Cell suspensions were sonicated on ice during 5 min at 35% amplitude. Cell lysates were centrifuged at 10,000 g at 4°C for 40 min. Supernates were loaded by gravity onto a Ni-^2+^-charged column (HisTrapHP, GE Healthcare) equilibrated with Buffer E. The column was washed with 20 mM imidazole in Buffer E until protein could not be detected by the Bradford protein assay, and his-FtsE was eluted by gradually increasing the concentration of in the mobile phase from 20 to 500 mM imidazole in Buffer E. Eluted proteins were collected in 1 ml fractions and examined by SDS-PAGE. Fractions with the highest purity were pooled and dialyzed overnight against 25 mM Tris-HCl pH 7.5, 2 mM DTT, and 100 mM NaCl at 4°C. The his-tag of the dialyzed protein was cleaved off by digestion with his-tagged TEV protease at 30°C for 3 hours. Undigested proteins, free his-tag and his-tagged TEV were separated from digested FtsE by performing a second immobilized metal affinity chromatography step with a Ni-^2+^-charged column. The digested FtsE protein was collected in the flow through and concentrated up to 8 mg/ml using a Millipore ultra-concentrator (10 kDa cutoff). Concentration of purified FtsE was determined by absorbance at 280 nm using an extinction coefficient of ε = 15,930 M^−1^ cm^−1^ calculated by the ExPASy ProtParam program (60).

### FtsE crystallization

Crystallization screenings were performed by high-throughput techniques in a NanoDrop robot and Innovadyne SD-2 microplates (Innovadyne Technologies Inc.), screening PACT Suite and JCSG Suite (Qiagen), JBScreen Classic 1-4 and 6 (Jena Bioscience) and Crystal Screen, Crystal Screen 2 and Index HT (Hampton Research). The conditions that produced crystals were optimized by sitting-drop vapor-diffusion method at 291 K by mixing 1 µL of protein solution and 1 µL of precipitant solution, equilibrated against 150 µL of precipitant solution in the reservoir chamber. The best crystals were obtained in a crystallization condition containing 0.15 M sodium acetate, 0.1 M Bis-Tris propane pH 6.5 and 16% (w/v) PEG3350 (Fig. S1B). Protein concentration was assayed at the concentration of 8 mg/mL.

### Soaking experiments with AMPPNP

For soaking trials, AMPPNP was dissolved in water and the pH was adjusted to 7.0 by addition of NaOH. Next, AMPPNP was incubated with native protein crystals at a final concentration of 5 mM using the crystallization conditions described above. Soaking experiments were incubated overnight at 291 K.

### Data collection, phasing and model refinement

Diffraction data was collected in beamlines ID23-1 at the European Synchrotron Radiation Facility (ESRF) and XALOC at the ALBA synchrotron, using the Pilatus 6M detector. Crystals diffracted up to 1.36-2.40 Å resolution and belonged to the P1 or P 2_1_ space groups, being the unit cell parameters *a* = 33.39 Å, *b* = 36.71 Å, *c* = 41.11 Å, α = 105.27°, γ = 95.69°, β = 99.69° and *a* = 33.11 Å, *b* = 118.79 Å, *c* = 82.48 Å, α = γ = 90°, β = 97.51°, respectively. The collected datasets were processed with XDS (61) and Aimless (62). In the P 1 crystals, one monomer was found in the asymmetric unit, yielding a Matthews coefficient (63) of 1.86 Å^3^/Da and a solvent content of 33.83%. In the P 2_1_ crystals, three monomers were found in the asymmetric unit, yielding a Matthews coefficient of 2.08 Å^3^/Da and a solvent content of 40.78%. For crystals belonging to space group P_1_, the structure determination was performed by *de novo* phasing with Arcimboldo (64). We used a search including 7 copies of a helix containing 6, 8, 10, 12, 13, 15 and 16 residues, respectively, assuming *rmsd* from target of 0.2 Å. For crystals belonging to space group P 2_1_ the structure determination was performed by the molecular replacement method. In both cases, the model was manually completed using Coot (65) followed by refinement using PHENIX (66). A summary of the refinement statistics is given in Table 1.

### Microscale Thermophoresis experiments

We ordered the synthetic sequence NTIRITIISREIQIMRLVGAKNSYIRG (JPT Peptide Technologies GmbH) for assaying the interaction with FtsE, but this peptide is extremely hydrophobic and could not been synthetized. Instead we used a shorter version, ISREIQIMRLVGA that could be successfully synthesized. For performing MST experiments, fluorescent label dye NT-647 was covalently attached to the protein. The concentration of labeled FtsE was kept constant (10 µM) while the concentration of non-labeled peptide was varied between 10 µM-0.3 nM. The assay was performed in buffer 25 mM Tris-HCl pH 7.5, 150 mM NaCl and 5 mM ATP. After 30 min incubation at RT the samples were loaded into Monolith™ NT.115 Premium Treated Capillaries and the MST analysis was performed using a Monolith NT.115. Unfortunately, we could not obtain any complex both by soaking or co-crystallization with the indicated peptide.

### Cultivation and transformation of S. pneumoniae

*S. pneumoniae* was grown in liquid C medium (67) or on Todd-Hewitt (BD Difco™) agar plates at 37°C. Plates were incubated in an anaerobic chamber made by including AnaeroGen™ bags from Oxoid in a sealed container. For selection of transformants, kanamycin or streptomycin were added to the growth medium to a final concentration of 400 and 200 µg/ml, respectively. Genetic transformation of *S. pneumoniae* was performed by natural competence as previously described by Straume *et al* (68).

### Construction of S. pneumoniae mutants

Construction of gene knockout cassettes and amplicons used to introduce genetic mutations in *S. pneumoniae* were done by using overlap extension PCR as previously described by Johnsborg *et al* (69). The primers used are listed in Table S2. Amplicons were flanked 5’ and 3’ with ∼1000 bp homologous with sequences flanking the target site in the *S. pneumoniae* genome allowing double cross-over via homologous recombination after transformation. Gene knockouts were confirmed by PCR, and mutations were verified by Sanger sequencing. The strains used in this work are listed in Table S1.

### Depletion of FtsX and microscopy

To deplete ectopic expression of FtsX in *S. pneumoniae* we used the ComRS gene expression/depletion system (70). For the growth experiment, strains ds312, ds314, ds751, ds753, ds754 and ds761 were grown to OD_550_ = 0.2 in five ml C medium containing 0.5 µM ComS inducer. Then the cells were collected at 4000 x g and washed once in five ml C medium without ComS. To start *ftsX* depletion, the cells were diluted to OD_550_ = 0.05 in C medium without ComS. The cultures were further diluted two-fold in ComS-free C medium in a series of 12 tubes and 300 µl from each dilution were transferred to a 96-well microtiter plate. The plate was incubated at 37°C and OD_550_ was measured automatically every fifth minute in a Synergy H1 Hybrid Reader (Biotek). For microscopic examinations of cells depleted of ectopically expressed FtsX, the strains ds312, ds314, ds751, ds753, ds754 and ds761 were pre-grown and washed as described above. Then the cultures were further diluted 128 times (corresponding to well eight in the microtiter plate experiment) in ComS-free C medium to a final volume of 10 ml. At OD_550_ = 0.2 or when the growth stopped at a lower density due to depletion of ectopic FtsX, cells were immobilized on a microscopy slide using 1.2% agarose. Images of the cells were taken by using a Zeiss AxioObserver with ZEN Blue software, and an ORCA-Flash 4.0 V2 Digital CMOS camera (Hamamatsu Photonics) with a 1003 phase-contrast objective. Images were prepared in ImageJ and cell detection for morphogenesis analyses were done by using the MicrobeJ plugin (71).

### Immunodetection of Flag-tagged FtsX

Flag-tagged versions of FtsX expressed in the native *ftsX* locus of strains ds768, ds769, ds770, ds771 and ds772 were detected by immunoblotting. The cells were grown in the presence of 0.05 µM ComS inducer to drive expression of ectopic FtsX. Cells from 10 ml cultures with OD_550_ = 0.2 were collected at 4000 x g, and resuspended in 200 µl SDS sample buffer. The samples were heated at 95°C for 10 minutes before proteins were separated by SDS-PAGE using a 12% separation gel. The proteins were then electroblotted onto a PVDF membrane (Biorad) and Flag-tagged FtsX proteins were detected by using ANTI-FLAG^®^ antibody (Sigma) as previously described by Stamsås *et al*. (72) except that both the primary and secondary anti rabbit-HRP antibodies (Sigma) were diluted 1:4000 in TBS-T.

## ACKNOWLEDGMENTS

We thank the staff from ALBA and ESRF synchrotron facilities for help during crystallographic data collection. This work was supported by grant number BFU2017-90030-P from the Spanish Ministry of Science, Innovation and Universities to J.A.H. The work was partly funded by the Research council of Norway.

## AUTHORS CONTRIBUTIONS

M. A., L.S.H. and J.A.H designed the experiments; M.A., D.S. performed the research; L.S.H., J.A.H., M.A., J.L., and D.S. analyzed the data; M.A., L.S.H. and J.A.H. wrote the paper. All authors discussed the results, edited and approved the paper.

## REFERENCES

1. Mavrici D, Marakalala MJ, Holton JM, Prigozhin DM, Gee CL, Zhang YJ, Rubin EJ, Alber T. 2014. Mycobacterium tuberculosis FtsX extracellular domain activates the peptidoglycan hydrolase, RipC. Proc Natl Acad Sci U S A 111:8037–8042.

2. Vollmer W. 2012. Bacterial growth does require peptidoglycan hydrolases. Mol Microbiol2012/10/17. 86:1031–1035.

3. Vollmer W, Joris B, Charlier P, Foster S. 2008. Bacterial peptidoglycan (murein) hydrolases. FEMS Microbiol Rev2008/02/13. 32:259–286.

4. Yang DC, Tan K, Joachimiak A, Bernhardt TG. 2012. A conformational switch controls cell wall-remodelling enzymes required for bacterial cell division. Mol Microbiol2012/06/22. 85:768–781.

5. Uehara T, Parzych KR, Dinh T, Bernhardt TG. 2010. Daughter cell separation is controlled by cytokinetic ring-activated cell wall hydrolysis. EMBO J.

6. Pichoff S, Du S, Lutkenhaus J. 2019. Roles of FtsEX in cell division. Res Microbiol.

7. Schmidt KL, Peterson ND, Kustusch RJ, Wissel MC, Graham B, Phillips GJ, Weiss DS. 2004. A predicted ABC transporter, FtsEX, is needed for cell division in Escherichia coli. J Bacteriol 186:785–793.

8. Du S, Pichoff S, Lutkenhaus J. 2016. FtsEX acts on FtsA to regulate divisome assembly and activity. Proc Natl Acad Sci U S A.

9. Arends SJ, Kustusch RJ, Weiss DS. 2009. ATP-binding site lesions in FtsE impair cell division. J Bacteriol2009/04/21. 191:3772–3784.

10. Yang DC, Peters NT, Parzych KR, Uehara T, Markovski M, Bernhardt TG. 2011. An ATP-binding cassette transporter-like complex governs cell-wall hydrolysis at the bacterial cytokinetic ring. Proc Natl Acad Sci 108:E1052–E1060.

11. Corbin BD, Wang Y, Beuria TK, Margolin W. 2007. Interaction between cell division proteins FtsE and FtsZ. J Bacteriol2007/02/20. 189:3026–3035.

12. Du S, Henke W, Pichoff S, Lutkenhaus J. 2019. How FtsEX localizes to the Z ring and interacts with FtsA to regulate cell division. Mol Microbiol2019/06/09.

13. Mir MA, Arumugam M, Mondal S, Rajeswari HS, Ramakumar S, Ajitkumar P. 2015. Mycobacterium tuberculosis cell division protein, FtsE, is an ATPase in dimeric form. Protein J2014/12/17. 34:35–47.

14. Locher KP. 2016. Mechanistic diversity in ATP-binding cassette (ABC) transporters. Nat Struct Mol Biol2016/06/09. 23:487–493.

15. Doige CA, Ames GF. 1993. ATP-dependent transport systems in bacteria and humans: relevance to cystic fibrosis and multidrug resistance. Annu Rev Microbiol 47:291–319.

16. Riordan JR, Rommens JM, Kerem B, Alon N, Rozmahel R, Grzelczak Z, Zielenski J, Lok S, Plavsic N, Chou JL, et al. 1989. Identification of the cystic fibrosis gene: cloning and characterization of complementary DNA. Science (80-)1989/09/08. 245:1066–1073.

17. Gottesman MM, Pastan I, Ambudkar S V. 1996. P-glycoprotein and multidrug resistance. Curr Opin Genet Dev1996/10/01. 6:610–617.

18. Greene NP, Kaplan E, Crow A, Koronakis V. 2018. Antibiotic Resistance Mediated by the MacB ABC Transporter Family: A Structural and Functional Perspective. Front Microbiol2018/06/13. 9:950.

19. Meisner J, Montero Llopis P, Sham LT, Garner E, Bernhardt TG, Rudner DZ. 2013. FtsEX is required for CwlO peptidoglycan hydrolase activity during cell wall elongation in Bacillus subtilis. Mol Microbiol 89:1069–1083.

20. Sham LT, Barendt SM, Kopecky KE, Winkler ME. 2011. Essential PcsB putative peptidoglycan hydrolase interacts with the essential FtsXSpn cell division protein in Streptococcus pneumoniae D39. Proc Natl Acad Sci U S A 108:E1061–9.

21. Bartual SG, Straume D, Stamsas GA, Munoz IG, Alfonso C, Martinez-Ripoll M, Havarstein LS, Hermoso JA. 2014. Structural basis of PcsB-mediated cell separation in Streptococcus pneumoniae. Nat Commun 5:3842.

22. Massidda O, Novakova L, Vollmer W. 2013. From models to pathogens: how much have we learned about Streptococcus pneumoniae cell division? Env Microbiol2013/07/16. 15:3133–3157.

23. Mesnage S, Chau F, Dubost L, Arthur M. 2008. Role of N-acetylglucosaminidase and N-acetylmuramidase activities in Enterococcus faecalis peptidoglycan metabolism. J Biol Chem2008/05/21. 283:19845–19853.

24. Sham LT, Jensen KR, Bruce KE, Winkler ME. 2013. Involvement of FtsE ATPase and FtsX extracellular loops 1 and 2 in FtsEX-PcsB complex function in cell division of Streptococcus pneumoniae D39. MBio2013/07/19. 4.

25. Ng WL, Kazmierczak KM, Winkler ME. 2004. Defective cell wall synthesis in Streptococcus pneumoniae R6 depleted for the essential PcsB putative murein hydrolase or the VicR (YycF) response regulator. Mol Microbiol2004/08/13. 53:1161–1175.

26. Ng WL, Robertson GT, Kazmierczak KM, Zhao J, Gilmour R, Winkler ME. 2003. Constitutive expression of PcsB suppresses the requirement for the essential VicR (YycF) response regulator in Streptococcus pneumoniae R6. Mol Microbiol2003/12/04. 50:1647–1663.

27. Giefing C, Meinke AL, Hanner M, Henics T, Bui MD, Gelbmann D, Lundberg U, Senn BM, Schunn M, Habel A, Henriques-Normark B, Ortqvist A, Kalin M, von Gabain A, Nagy E. 2008. Discovery of a novel class of highly conserved vaccine antigens using genomic scale antigenic fingerprinting of pneumococcus with human antibodies. J Exp Med2008/01/02. 205:117–131.

28. Giefing-Kroll C, Jelencsics KE, Reipert S, Nagy E. 2011. Absence of pneumococcal PcsB is associated with overexpression of LysM domain-containing proteins. Microbiology2011/04/09. 157:1897–1909.

29. Kempf MJ, McBride MJ. 2000. Transposon insertions in the Flavobacterium johnsoniae ftsX gene disrupt gliding motility and cell division. J Bacteriol2000/02/29. 182:1671–1679.

30. Merino S, Altarriba M, Gavin R, Izquierdo L, Tomas JM. 2001. The cell division genes (ftsE and X) of Aeromonas hydrophila and their relationship with opsonophagocytosis. FEMS Microbiol Lett2001/06/30. 198:183–188.

31. Ramirez-Arcos S, Salimnia H, Bergevin I, Paradis M, Dillon JA. 2001. Expression of Neisseria gonorrhoeae cell division genes ftsZ, ftsE and minD is influenced by environmental conditions. Res Microbiol2002/01/05. 152:781–791.

32. Salmond GP, Plakidou S. 1984. Genetic analysis of essential genes in the ftsE region of the Escherichia coli genetic map and identification of a new cell division gene, ftsS. Mol Gen Genet1984/01/01. 197:304–308.

33. Garti-Levi S, Hazan R, Kain J, Fujita M, Ben-Yehuda S. 2008. The FtsEX ABC transporter directs cellular differentiation in Bacillus subtilis. Mol Microbiol2008/06/25. 69:1018–1028.

34. Meier EL, Daitch AK, Yao Q, Bhargava A, Jensen GJ, Goley ED. 2017. FtsEX-mediated regulation of the final stages of cell division reveals morphogenetic plasticity in Caulobacter crescentus. PLoS Genet2017/09/09. 13:e1006999.

35. Li Q, Li Z, Li X, Xia L, Zhou X, Xu Z, Shao J, Shen Q, Zhang R. 2018. FtsEX-CwlO regulates biofilm formation by a plant-beneficial rhizobacterium Bacillus velezensis SQR9. Res Microbiol2018/02/11. 169:166–176.

36. Hung LW, Wang IX, Nikaido K, Liu PQ, Ames GF, Kim SH. 1998. Crystal structure of the ATP-binding subunit of an ABC transporter. Nature 396:703–707.

37. Ye J, Osborne AR, Groll M, Rapoport TA. 2004. RecA-like motor ATPases--lessons from structures. Biochim Biophys Acta2004/10/30. 1659:1–18.

38. Rees DC, Johnson E, Lewinson O. 2009. ABC transporters: the power to change. Nat Rev Mol Cell Biol 10:218–227.

39. Finn RD, Mistry J, Tate J, Coggill P, Heger A, Pollington JE, Gavin OL, Gunasekaran P, Ceric G, Forslund K, Holm L, Sonnhammer EL, Eddy SR, Bateman A. 2010. The Pfam protein families database. Nucleic Acids Res2009/11/19. 38:D211–22.

40. Pai EF, Kabsch W, Krengel U, Holmes KC, John J, Wittinghofer A. 1989. Structure of the guanine-nucleotide-binding domain of the Ha-ras oncogene product p21 in the triphosphate conformation. Nature 341:209–214.

41. Abrahams JP, Leslie AG, Lutter R, Walker JE. 1994. Structure at 2.8 A resolution of F1-ATPase from bovine heart mitochondria. Nature 370:621–628.

42. Laskowski RA, Swindells MB. 2011. LigPlot+: multiple ligand-protein interaction diagrams for drug discovery. J Chem Inf Model2011/09/17. 51:2778–2786.

43. Okada U, Yamashita E, Neuberger A, Morimoto M, van Veen HW, Murakami S. 2017. Crystal structure of tripartite-type ABC transporter MacB from Acinetobacter baumannii. Nat Commun2017/11/08. 8:1336.

44. Crow A, Greene NP, Kaplan E, Koronakis V. 2017. Structure and mechanotransmission mechanism of the MacB ABC transporter superfamily. Proc Natl Acad Sci U S A2017/11/08. 114:12572–12577.

45. Chen J, Lu G, Lin J, Davidson AL, Quiocho FA. 2003. A tweezers-like motion of the ATP-binding cassette dimer in an ABC transport cycle. Mol Cell2003/10/07. 12:651–661.

46. Rued BE, Alcorlo M, Edmonds KA, Martinez-Caballero S, Straume D, Fu Y, Bruce KE, Wu H, Havarstein LS, Hermoso JA, Winkler ME, Giedroc DP. 2019. Structure of the Large Extracellular Loop of FtsX and Its Interaction with the Essential Peptidoglycan Hydrolase PcsB in Streptococcus pneumoniae. MBio2019/01/31. 10.

47. Waterhouse A, Bertoni M, Bienert S, Studer G, Tauriello G, Gumienny R, Heer FT, de Beer TAP, Rempfer C, Bordoli L, Lepore R, Schwede T. 2018. SWISS-MODEL: homology modelling of protein structures and complexes. Nucleic Acids Res2018/05/23. 46:W296–W303.

48. Fitzpatrick AWP, Llabres S, Neuberger A, Blaza JN, Bai XC, Okada U, Murakami S, van Veen HW, Zachariae U, Scheres SHW, Luisi BF, D. D. 2017. Structure of the MacAB-TolC ABC-type tripartite multidrug efflux pump. Nat Microbiol2017/05/16. 2:17070.

49. Lee JY, Kinch LN, Borek DM, Wang J, Wang J, Urbatsch IL, Xie XS, Grishin N V, Cohen JC, Otwinowski Z, Hobbs HH, Rosenbaum DM. 2016. Crystal structure of the human sterol transporter ABCG5/ABCG8. Nature2016/05/05. 533:561–564.

50. Bouige P, Laurent D, Piloyan L, Dassa E. 2005. Phylogenetic and Functional Classification of ATP-Binding Cassette (ABC) Systems. Curr Protein Pept Sci.

51. Daley DO, Rapp M, Granseth E, Melén K, Drew D, Von Heijne G. 2005. Biochemistry: Global topology analysis of the Escherichia coli inner membrane proteome. Science (80-).

52. Kobayashi N, Nishino K, Hirata T, Yamaguchi A. 2003. Membrane topology of ABC-type macrolide antibiotic exporter MacB in Escherichia coli. FEBS Lett.

53. Tokuda H, Matsuyama SI. 2004. Sorting of lipoproteins to the outer membrane in E. coli. Biochim Biophys Acta - Mol Cell Res.

54. Yakushi T, Masuda K, Narita SI, Matsuyama SI, Tokuda H. 2000. A new ABC transporter mediating the detachment of lipid-modified proteins from membranes. Nat Cell Biol.

55. Yamanaka H, Kobayashi H, Takahashi E, Okamoto K. 2008. MacAB is involved in the secretion of Escherichia coli heat-stable enterotoxin II. J Bacteriol.

56. Egan AJF, Vollmer W. 2013. The physiology of bacterial cell division. Ann N Y Acad Sci 1277:8–28.

57. Brunet YR, Wang X, Rudner DZ. 2019. SweC and SweD are essential co-factors of the FtsEX-CwlO cell wall hydrolase complex in Bacillus subtilis. PLoS Genet.

58. Straume D, Piechowiak KW, Olsen S, Stamsås GA, Berg KH, Kjos M, Heggenhougen MV, Alcorlo M, Hermoso JA, Håvarstein LS. 2020. Class A PBPs have a distinct and unique role in the construction of the pneumococcal cell wall. Proc Natl Acad Sci U S A.

59. Yang DC, Peters NT, Parzych KR, Uehara T, Markovski M, Bernhardt TG. 2011. An ATP-binding cassette transporter-like complex governs cell-wall hydrolysis at the bacterial cytokinetic ring. Proc Natl Acad Sci U S A.

60. Gasteiger E, Gattiker A, Hoogland C, Ivanyi I, Appel RD, Bairoch A. 2003. ExPASy: The proteomics server for in-depth protein knowledge and analysis. Nucleic Acids Res2003/06/26. 31:3784–3788.

61. Kabsch W. 2010. Xds. Acta Crystallogr D Biol Crystallogr 66:125–132.

62. Evans PR, Murshudov GN. 2013. How good are my data and what is the resolution? Acta Crystallogr D Biol Crystallogr 69:1204–1214.

63. Matthews BW. 1968. Solvent content of protein crystals. J Mol Biol 33:491–497.

64. Rodriguez D, Sammito M, Meindl K, de Ilarduya IM, Potratz M, Sheldrick GM, Uson I. 2012. Practical structure solution with ARCIMBOLDO. Acta Crystallogr D Biol Crystallogr 68:336–343.

65. Emsley P, Lohkamp B, Scott WG, Cowtan K. 2010. Features and development of Coot. Acta Crystallogr D Biol Crystallogr 66:486–501.

66. Adams PD, Afonine P V, Bunkoczi G, Chen VB, Davis IW, Echols N, Headd JJ, Hung LW, Kapral GJ, Grosse-Kunstleve RW, McCoy AJ, Moriarty NW, Oeffner R, Read RJ, Richardson DC, Richardson JS, Terwilliger TC, Zwart PH. 2010. PHENIX: a comprehensive Python-based system for macromolecular structure solution. Acta Crystallogr D Biol Crystallogr 66:213–221.

67. Lacks S, Hotchkiss RD. 1960. A study of the genetic material determining an enzyme activity in pneumococcus. Biochim Biophys Acta 39:508–518.

68. Straume D, Stamsås GA, Berg KH, Salehian Z, Håvarstein LS. 2017. Identification of pneumococcal proteins that are functionally linked to penicillin-binding protein 2b (PBP2b). Mol Microbiol2016/10/28. 103:99–116.

69. Johnsborg O, Eldholm V, Bjørnstad ML, Håvarstein LS. 2008. A predatory mechanism dramatically increases the efficiency of lateral gene transfer in Streptococcus pneumoniae and related commensal species. Mol Microbiol2008/05/20. 69:245–253.

70. Berg KH, Biørnstad TJ, Straume D, Håvarstein LS. 2011. Peptide-regulated gene depletion system developed for use in Streptococcus pneumoniae. J Bacteriol2011/08/02. 193:5207–5215.

71. Ducret A, Quardokus EM, Brun Y V. 2016. MicrobeJ, a tool for high throughput bacterial cell detection and quantitative analysis. Nat Microbiol.

72. Stamsås GA, Straume D, Salehian Z, Håvarstein LS. 2017. Evidence that pneumococcal WalK is regulated by StkP through protein-protein interaction. Microbiology2016/12/03. 163:383–399.

